# Single-Site Phosphorylation Elicits Structural, Dynamic, and Accessibility Changes in Proteins at both Proximal and Distal Regions to the Phosphosite

**DOI:** 10.1101/2023.11.30.569386

**Authors:** Seemadri Subhadarshini, Himani Tandon, Narayanaswamy Srinivasan, Ramanathan Sowdhamini

**Author notes:** This article is dedicated to the memory of late Prof. Narayanaswamy Srinivasan.

## Abstract

Phosphorylation, a fundamental cellular mechanism, intricately regulates protein function and signaling pathways. Our study employs extensive computational analyses on a curated dataset of phosphorylated and unphosphorylated protein structures to explore the multifaceted impact of phosphorylation on protein conformation. Our findings reveal that phosphorylation induces not only local changes at the phosphorylation site but also extensive alterations in distant regions, showcasing its far-reaching influence on protein structure-dynamics. Using Normal Mode Analysis (NMA), we investigate changes in protein flexibility post-phosphorylation, highlighting an enhanced level of structural dynamism. Through in-depth case studies on Polyubiquitin-B and Glycogen Synthase Kinase-3 Beta, we elucidate how phosphorylation at distinct sites leads to variable structural and dynamic modifications, potentially dictating functional outcomes. While phosphorylation largely preserves residue motion correlation, it significantly disrupts low-frequency global modes, presenting a dualistic impact on protein dynamics. We also explore alterations in the total accessible surface area (ASA), emphasizing region-specific changes around phosphorylation sites. This study sheds light on phosphorylation-induced conformational changes, dynamic modulation, and surface accessibility alterations, contributing to a comprehensive understanding of cellular regulation and suggesting promising avenues for therapeutic interventions.

## INTRODUCTION

Phosphorylation is a prevalent and crucial post-translational modification (PTM) that plays a fundamental role in regulating protein function and cellular signaling. It is involved in a wide range of cellular processes, including cell cycle regulation, metabolism, signal transduction, and gene expression [1, 2]. In eukaryotes, serine, threonine, and tyrosine residues are frequently phosphorylated. Among these, serine is the most predominant, followed by threonine and then tyrosine [3]. On the other hand, in prokaryotes, histidine residues serve as the primary phosphorylation sites. This enzymatic addition of a phosphate group, facilitated by protein kinases, acts as a molecular switch, modulating protein activity, localization, and interactions [4]. Phosphorylation is a highly regulated process that enables cells to rapidly respond to external stimuli by activating or deactivating specific proteins or signaling cascades [5]. It exerts its regulatory effects through various mechanisms, including inducing conformational changes that expose or hide functional domains, creating binding sites for other proteins or disrupting existing interactions, influencing subcellular localization, and regulating enzymatic activity. These phosphorylation events orchestrate intricate cellular signaling networks [6, 7], enabling cells to adapt and respond to changing environmental conditions.

Numerous studies have demonstrated that when proteins interact with effector molecules, such as small molecules, proteins, DNA/RNA, mutations, or other post-translational modifications [8], they undergo structural modifications, changes in dynamics and flexibility, and alterations in surface accessibility. These changes can occur not only at the binding site but also in regions distant from it. The effector molecule can induce significant conformational alterations in the protein, either locally or globally, or cause subtle shifts in the equilibrium between different conformations [8, 9]. The present study focuses on elucidating the intricacies of phosphorylation-induced structure-dynamics-accessibility alterations. By focussing on this post-translational modification, our research aims to contribute to a more comprehensive understanding of the consequential change in the conformational landscape of proteins, associated with the phosphorylation events.

Phosphorylation-induced conformational changes refer to alterations in the three-dimensional structure of proteins upon the addition of a phosphate moiety. These changes can impact the overall folding of the protein or specific regions, leading to modifications in its functional properties. For instance, phosphorylation may induce conformational shifts that expose or conceal binding sites, influencing the protein’s ability to interact with other molecules such as substrates, cofactors, or other proteins in a signaling cascade [10]. Phosphorylation not only affects static conformation but also introduces dynamic changes in protein behaviour [11]. Proteins are not rigid structures; rather, they exhibit inherent flexibility and dynamic motion. Phosphorylation can influence these dynamic properties by altering the vibrational modes, flexibility, and motion within the protein structure. This dynamic modulation is essential for the fine-tuning of cellular processes, allowing for rapid and precise responses to various stimuli. Understanding how phosphorylation impacts protein dynamics provides insights into the temporal aspects of cellular signaling and regulatory mechanisms. Phosphorylation-induced changes extend beyond the protein’s core structure to its surface properties, specifically alterations in the accessible surface area (ASA). The addition of a phosphate group can expose or shield specific regions on the protein surface, influencing its interactions with solvent molecules, other proteins, or cellular membranes. These ASA changes are crucial for mediating protein-protein interactions, subcellular localization, and interactions with ligands [12]. By modulating the ASA, phosphorylation contributes to the spatiotemporal regulation of cellular events and ensures the precise orchestration of signaling cascades. The study also underscores the centrality of eukaryotic phosphorylation on serine, threonine, and tyrosine residues as a fundamental mechanism for regulating diverse cellular processes. Each type of phosphorylation, coupled with its distinct location on the amino acid residues, carries specific implications for protein structure, function, and cellular signaling. This nuanced understanding is paramount for unravelling the complexities of cellular regulation and holds the key to developing targeted therapeutic strategies.

Our current study takes a systematic approach to explore the prevalence, extent, location, and functional relevance of conformational alterations induced by phosphorylation at a single residue position. Analysing a non-redundant dataset of 24 protein pairs in both phosphorylated and unphosphorylated states, our study showcases distinct conformational changes occurring away from the phosphorylation site. Furthermore, we shed light on dynamic shifts in protein residues, emphasizing the long-range impact of phosphorylation on protein dynamics. The investigation also delves into how phosphorylation affects residue motion correlation and perturbs low-frequency global modes in the unphosphorylated form. Notably, observable differences in the total accessible surface post-phosphorylation are accentuated in the vicinity of the phosphorylation site, underlining the localized impact on surface properties. To provide a more granular understanding, the study delves into two case studies spanning serine, threonine, and tyrosine phosphorylation. The first case demonstrates how phosphorylation at different protein sites induces variable structural and dynamic changes, potentially impacting function. In the second case, we elucidate how phosphorylation within a kinase’s activation loop brings about functional changes by influencing its structure and dynamics. Collectively, our results strongly suggest that phosphorylation at a single residue is sufficient to cause alterations along the structure-dynamics-accessibility axes, emphasizing the pivotal role of this PTM in shaping cellular events and signaling cascades.

## RESULTS

### Protein structures undergo distinct conformational changes upon phosphorylation, with more pronounced alternations occurring away from the site of phosphorylation

SSP (Single-Site Phosphorylation) dataset, comprising of phosphorylated and unphosphorylated states of proteins, phosphorylated at a single Serine, Threonine or Tyrosine residue was specifically curated following the workflow outlined in **Supplementary Figure 1A** (refer Methods). In the SSP dataset, the phosphate moiety or modified protein residue showed a predominant presence in the loop region (50%), followed by the helix region (42%), with only a small percentage (8%) found in the beta-sheet region (**Supplementary Figure 1B**). Post dataset curation, the Cα-Root Mean Square Deviation (RMSD) between phosphorylated and unphosphorylated states was computed for each protein pair in the SSP dataset. Notably, 18 out of 24 proteins (∼75% of the dataset) exhibited significant structural differences (**Figure 1A**). Significance was determined by considering RMSD values exceeding the standard deviation from the mean of RMSDs observed in the control dataset. All protein pairs had a TM score value, higher than 0.5 indicating overall fold similarity (**Supplementary Figure 2A**). We compared the Cα-RMSD distribution of the SSP dataset with that of the control dataset in order to take into account the impact of crystal packing on the protein structure. The two distributions were found to be significantly different (two-sample Kilmogorov-Smirnov [KS] test and T-test, p < 10^-2^) indicating that phosphorylation modification of the protein, rather than crystallisation artefacts, is the primary cause of the observed changes in global protein conformation (**Figure 1B, C**).

**Figure 1:**
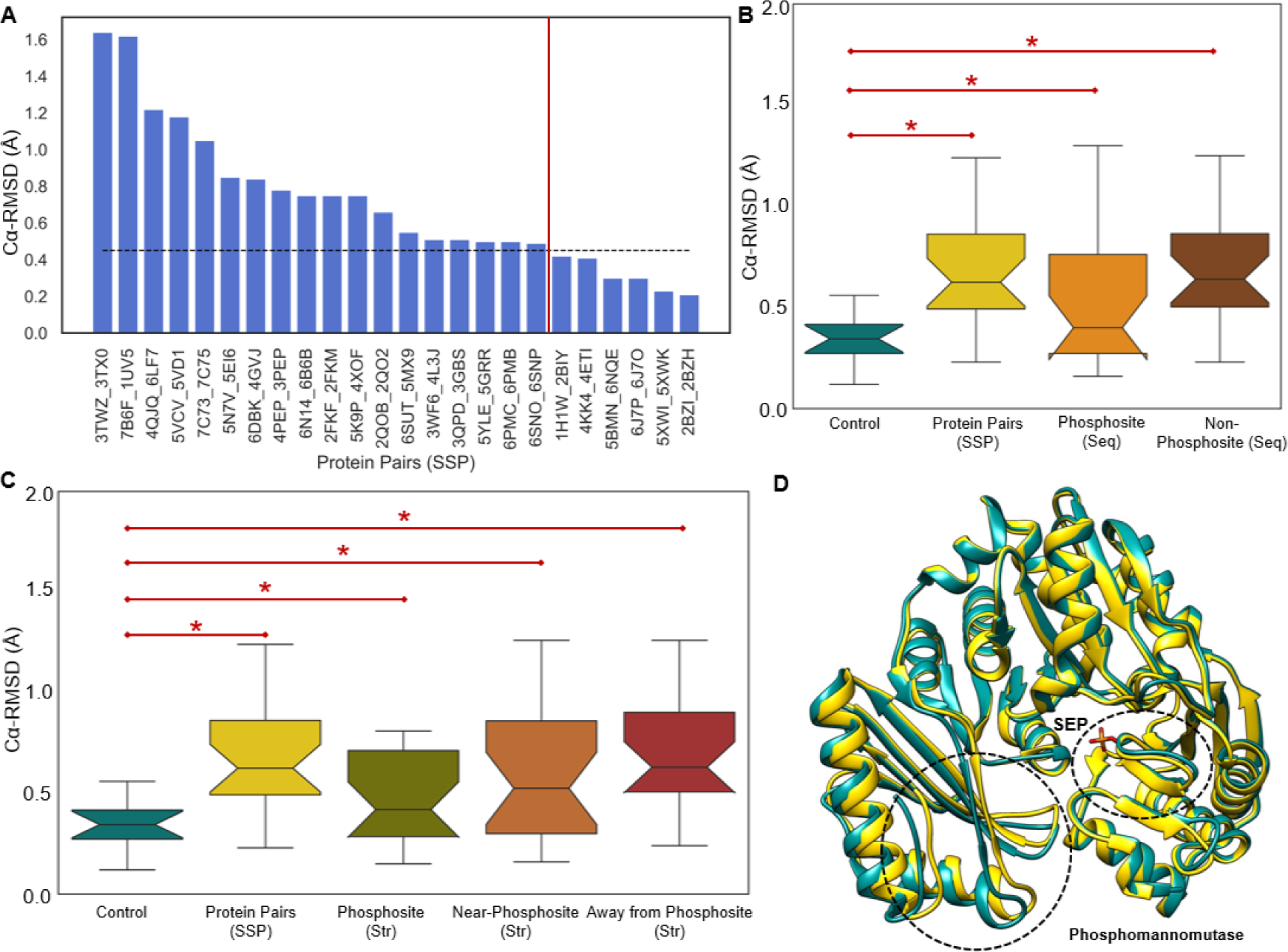
Comparative Structural Analyses of SSP Dataset. **(A)** Bar plot depicting Cα-RMSD for protein pairs. A higher RMSD indicates greater structural disparity between phosphorylated and unphosphorylated forms. The black line denotes the cut-off derived from the control dataset, with bars to the left of the red line signifying significant structural changes and those to the right as not significant. **(B)** Box plots showing the distribution of Cα-RMSD (Å) for the Control dataset, Protein Pairs in the SSP dataset, Phosphosite (Seq), and Non-Phosphosite (Seq). **(C)** Box plots displaying the distribution of Cα-RMSD (Å) for the Control dataset, Protein Pairs in the SSP dataset, Phosphosite (Str), Near-Phosphosite (Str), and Away from Phosphosite (Str). * signifies significant differences in distributions (two-sample KS test and T-test, p-value < 0.01). **(D)** Example case of Phosphomannomutase, presenting superimposed and aligned structures of the phosphorylated (PDB ID: 2FKF, coloured dark cyan) and unphosphorylated (PDB ID: 2FKM, coloured yellow) forms. Dotted circles highlight notable structural changes at the phosphosite and away from it.

We categorised the residues into *Phosphosite (Seq), Non-Phosphosite (Seq), Phosphosite (Str), Near-Phosphosite (Str),* and *Away from Phosphosite (Str)* residues, to identify specific regions undergoing conformational alterations linked to the phosphorylation event (refer Methods). RMSD was computed independently for these residue segments. This allowed us to comprehensively delineate the local structural differences between the phosphorylated and unphosphorylated states. The distribution of RMSD for both *Phosphosite (Seq)* and *Non-Phosphosite (Seq)* residues exhibited a statistically significant difference (two-sample Kilmogorov-Smirnov [KS] test and T-test, p < 10^-2^) compared to the control dataset (**Figure 1B**). This distinction was also observed in the case of *Phosphosite (Str), Near-Phosphosite (Str),* and *Away from Phosphosite (Str)* with respect to the control dataset (**Figure 1C**). Upon careful analysis of the plots (**Figure 1B, C**, and **Supplementary Figure 2B, C**, which includes outliers), it was observed that there are considerable structural differences between the phosphorylated and unphosphorylated forms at the phosphosite. Additionally, significant RMSD variations were found in the regions surrounding the phosphorylation site, both proximal and distal, with the disparities being more prominent in the latter case. A specific instance of the protein Phosphomannomutase, representing superimposed and aligned structures of both the phosphorylated (PDB ID: 2FKF) and unphosphorylated (PDB ID: 2FKM) forms, highlights notable structural changes at the phosphosite and in regions distal to it (**Figure 1D**).

### Phosphorylation induces dynamic shifts in protein residues, with notable effects extending further from the phosphorylation site, highlighting its long-range impact on protein dynamics

Protein flexibilities are a result of energetically feasible intrinsic functional motions encoded within their tertiary structures. These intrinsic motions, crucial for understanding the dynamic behaviour of biomolecules, are often investigated using Normal Mode Analysis (NMA) [9, 13]. By studying the vibrational motion of a harmonic oscillating system in the immediate vicinity of its equilibrium, NMA provides insights into the inherent structural dynamics of proteins. The structural flexibility of proteins, evident in their normal modes, plays a pivotal role in facilitating functionally important conformational variations [14, 15]. In order to explore the nuanced differences in residue dynamics between phosphorylated and unphosphorylated forms of the proteins, we conducted detailed analyses on the normalised square fluctuations obtained through Anisotropic Network Model based Normal Mode Analysis (ANM-NMA) on 34 (17*2) protein structures within the SSP dataset, after filtering out and excluding cases with missing residues in either the phosphorylated or unphosphorylated forms of the proteins (refer Methods).

The difference in distribution of absolute difference in normalised square fluctuations between the phosphorylated and unphosphorylated forms, was found to be statistically significant (two-sample KS test, p-value < 10^-2^), compared to the control group (**Figure 2A** and **Supplementary Figure 4A**, which includes outliers). Additionally, the distribution of normalized square fluctuation across all residues of phosphorylated proteins was significantly different (two-sample KS test, p-value < 10^-2^) from that of unphosphorylated proteins (**Figure 2B** and **Supplementary Figure 4B**, which includes outliers). This suggests that phosphorylated proteins exhibit altered flexibility in numerous residues post-phosphorylation, as indicated by a greater variance in fluctuation distribution. Upon further scrutiny, it was observed that, majority of the phosphorylated proteins exhibited a modestly higher root mean square fluctuation (RMSF) compared to their unphosphorylated counterparts (**Figure 2C**). This trend persisted even at a residue level, with a general increase in flexibility among residues following phosphorylation (**Figure 2D**). To quantify the percentage of residues undergoing significant changes, the difference in residue fluctuations between phosphorylated and unphosphorylated forms was calculated. Significance was attributed only if the difference exceeded the standard deviation from the mean of the fluctuation difference in the control dataset. The findings indicated that 3.4% of residues exhibited significantly higher fluctuation in the unphosphorylated form, while 1.4% of residues showed significantly higher fluctuation in the phosphorylated form. To ensure the robustness of these differences, irrespective of the 15 Å distance cutoff used for normal mode analysis (NMA) calculations, additional analyses were conducted. Normalized fluctuations of phosphorylated and unphosphorylated proteins in the SSP dataset, as well as all proteins in the control dataset, were calculated at 12 Å and 10 Å cutoffs. The absolute difference in fluctuations between the control and SSP dataset (**Supplementary Figure 3A**), as well as the fluctuations in phosphorylated and unphosphorylated forms (**Supplementary Figure 3B**), remained consistently significant at 12 Å and 10 Å cutoffs, indicating that these observed variations were insensitive to the chosen distance cutoff. These observations highlight a discernible post-phosphorylation augmentation in the dynamic motion of residues, signifying an enhanced level of structural flexibility.

**Figure 2:**
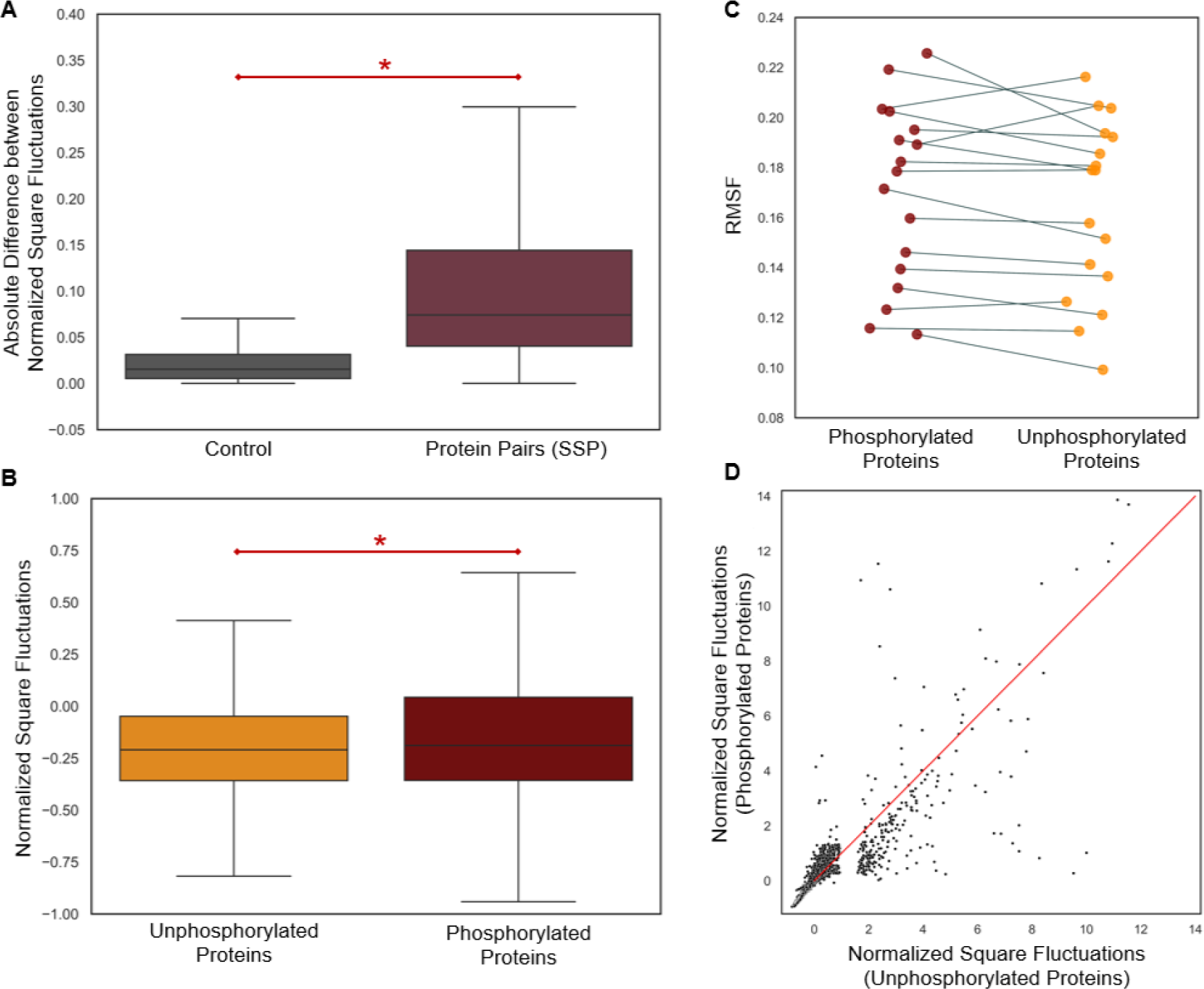
Dynamic Shifts. **(A)** Box plots illustrating the distribution of the absolute difference in normalized square fluctuation of proteins between the control dataset and the phosphorylated/unphosphorylated forms in the SSP dataset. **(B)** Paired strip plot of RMSF values for corresponding phosphorylated and unphosphorylated proteins in SSP dataset. **(C)** Box plot displaying the distribution of normalized square fluctuations for all residues in phosphorylated and unphosphorylated proteins. * indicates significant differences (two-sample KS test, p < 0.01) **(D)** Scatterplot presenting the fluctuations for all residues in phosphorylated and unphosphorylated forms, aiding in the identification of residues exhibiting increased or decreased fluctuations.

The fluctuation profiles of phosphorylated and unphosphorylated proteins were subject to separate analyses across distinct residue categories: *Phosphosite (Seq)*, *Non-Phosphosite (Seq)*, *Phosphosite (Str)*, *Near-Phosphosite (Str)*, and *Away from Phosphosite (Str)* (**Figure 3**). The objective was to pinpoint specific areas undergoing dynamic shifts associated with the phosphorylation modification event. In all regions-*Phosphosite (Seq)* (**Figure 3A**), *Non-Phosphosite (Seq)* (**Figure 3B**), *Phosphosite (Str)* (**Figure 3C**), *Near-Phosphosite (Str)* (**Figure 3D**), and *Away from Phosphosite (Str)* (**Figure 3E**)-the distributions of normalised square fluctuations of phosphorylated proteins were found to be significantly different (two-sample KS test, p-value < 10^-2^) compared to their unphosphorylated counterparts (**Supplementary Figure 4C,D**, which includes outliers). Notably, phosphorylated proteins exhibited marginally heightened level of flexibility [11] across all the aforementioned regions, with a particularly noteworthy observation near the phosphorylation sites. In this specific region *Near-Phosphosite (Str),* 1.1% of residues exhibited significantly higher fluctuation in the phosphorylated proteins, compared to only 0.2% in their unphosphorylated counterparts. These observation underscores the nuanced and localized impact of phosphorylation on the dynamic behaviour of proteins, across diverse residue contexts.

**Figure 3:**
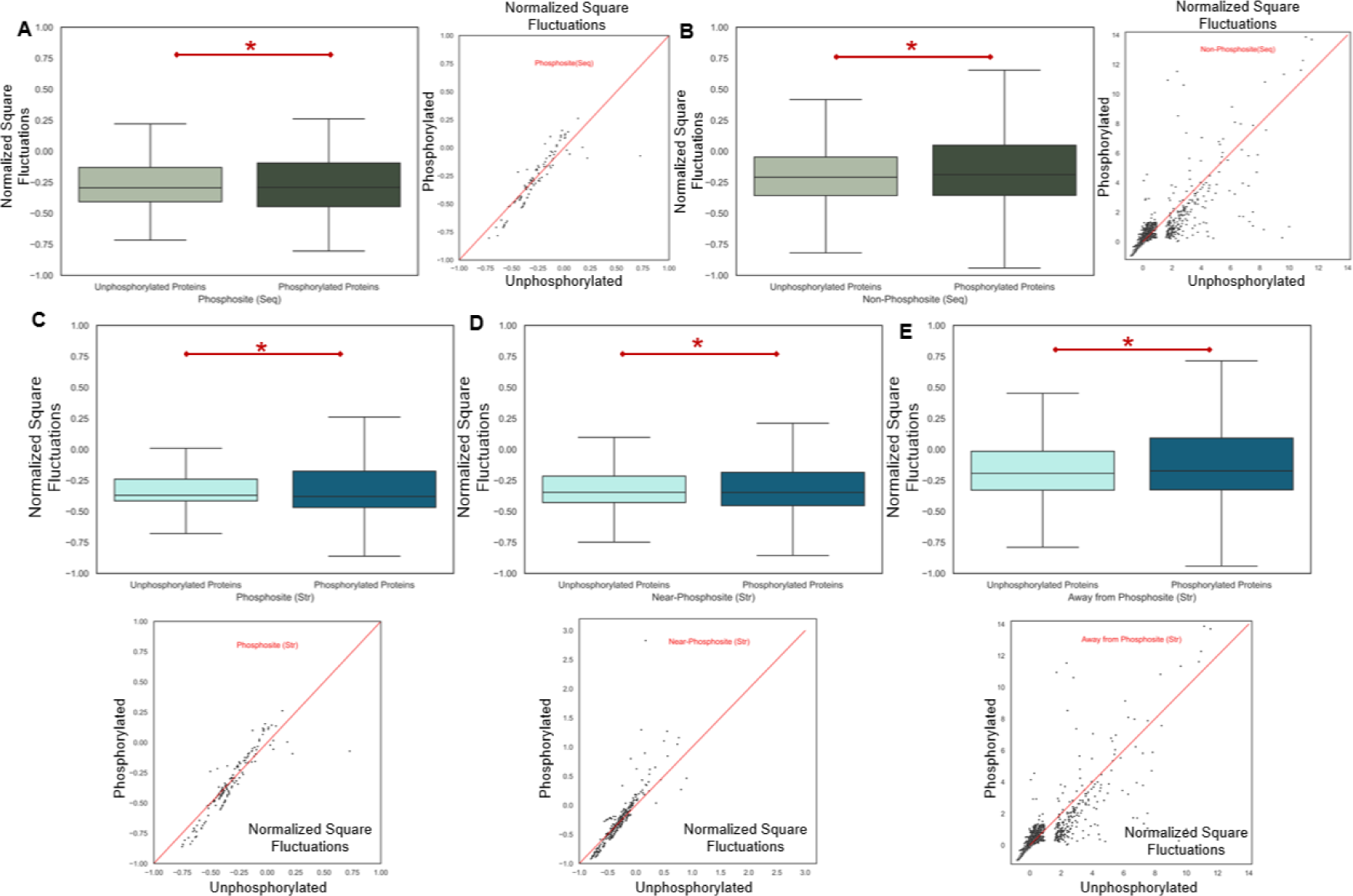
Localized Regions Showing Changes in Flexibility/Dynamics Post Phosphorylation. **(A)** Box plot (left) and scatterplot (right) showing normalized square fluctuations for *Phosphosite (Seq)* residues in phosphorylated and unphosphorylated proteins. **(B)** Box plot (left) and scatterplot (right) illustrating normalized square fluctuations for *Non-Phosphosite (Seq)* residues in phosphorylated and unphosphorylated proteins. **(C)** Box plot (top) and scatterplot (bottom) presenting normalized square fluctuations for *Phosphosite (Str)* residues in phosphorylated and unphosphorylated proteins. **(D)** Box plot (top) and scatterplot (bottom) displaying normalized square fluctuations for *Near-Phosphosite (Str)* residues in phosphorylated and unphosphorylated proteins. **(E)** Box plot (top) and scatterplot (bottom) depicting normalized square fluctuations for *Away from Phosphosite (Str)* residues in phosphorylated and unphosphorylated proteins.

### Protein phosphorylation does not significantly change residue motion correlation, but it does perturb the low-frequency global modes in the unphosphorylated form

Exploring the repercussions of protein phosphorylation on residue motion correlation is crucial for gaining insights into the complexities of residue communication within a protein. Correlated fluctuations among residues serve as conduits for vital information transmission, and identifying residues with coupled motion is instrumental in deciphering functional pathways [16]. Our study delved into the question of whether phosphorylation induces alterations in these dynamic interactions. Residue-residue cross-correlation matrices were computed for both phosphorylated and unphosphorylated forms of proteins in the SSP dataset, for cases where ANM-NMA analyses were performed. The Rv coefficient, calculated between the cross-correlation matrices of phosphorylated and unphosphorylated protein pairs, served as a quantitative measure. The coupling between residue fluctuations appeared to be largely unaffected post-phosphorylation, as indicated by high R_v_ coefficients (≥0.7) (**Figure 4A**). **Figure 4B** illustrates an example from the dataset featuring the protein Phosphomannomutase, where phosphorylation has no significant impact on the residue-residue correlation, as evidenced by a high R_v_ coefficient of 0.9. The conserved correlation patterns suggest that, despite undergoing phosphorylation, the fundamental communication pathways and cooperative interactions among residues remained largely unperturbed. This stable network of correlated fluctuations implies a robust and finely tuned system, where post-translational modifications such as phosphorylation at a single residue may not necessarily disrupt the intricate interplay between residues.

**Figure 4:**
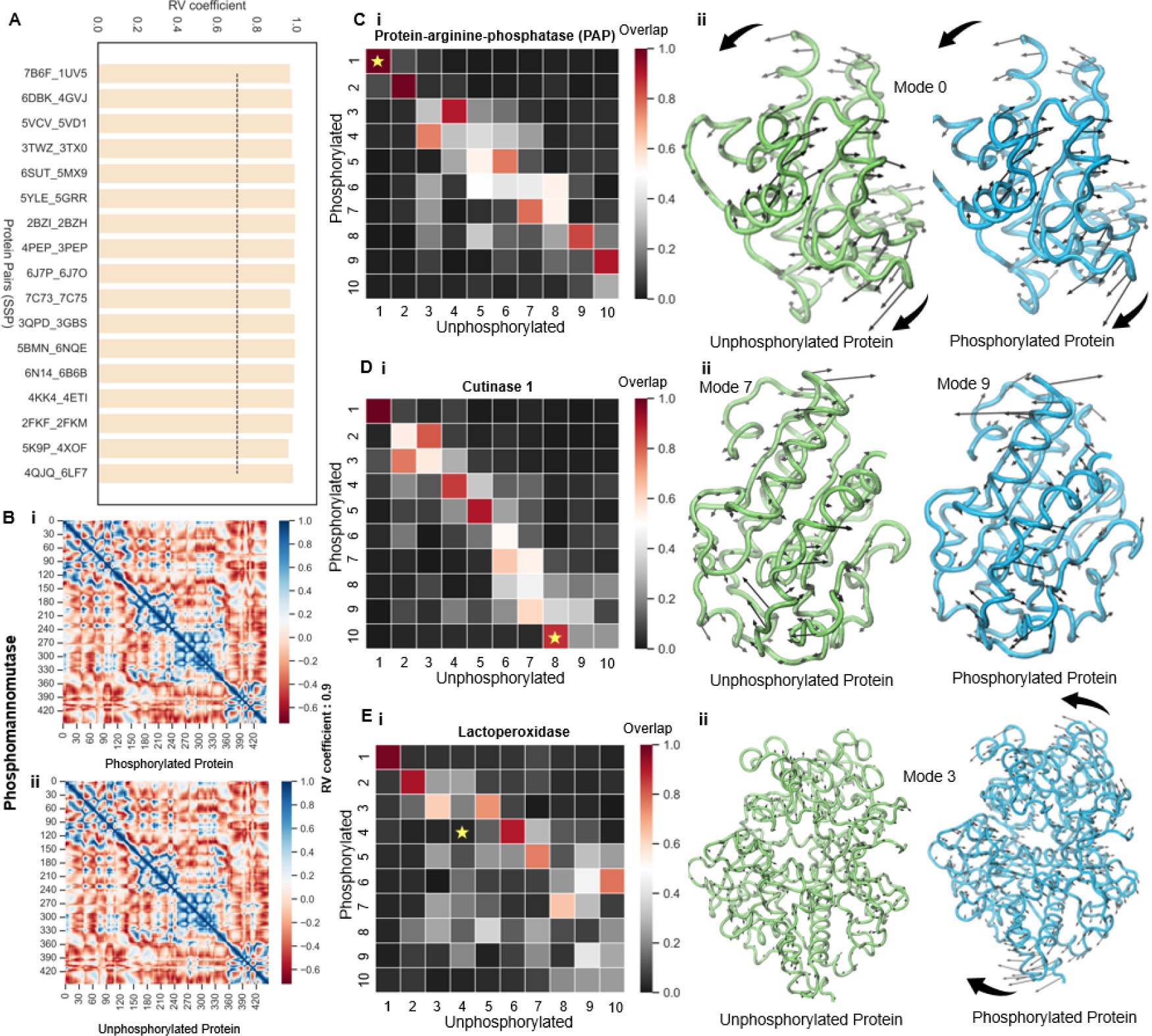
Cross-correlation Analysis and Overlap Analysis for Proteins in the SSP Dataset. **(A)** Bar plot representing the similarity between cross-correlation matrices of phosphorylated and unphosphorylated proteins. The black horizontal line denotes the similarity cutoff. **(B)** Example case from the SSP dataset featuring Phosphomannomutase (PDB ID: 2FKF, phosphorylated and PDB ID: 2FKM, unphosphorylated) where phosphorylation does not alter residue communication. The cross-correlation matrices show a high similarity (R_v_ coefficient= 0.9). **(C) (i)** Overlap of the first ten non-zero normal modes of phosphorylated and unphosphorylated forms of Protein-arginine-phosphatase (PAP). **(ii)** Backbone fluctuations indicated by eigen vectors of the unphosphorylated protein (left) and phosphorylated protein (right) for mode 1. **(D) (i)** Overlap of the first ten non-zero normal modes of phosphorylated and unphosphorylated forms of Cutinase1. **(ii)** Backbone fluctuations indicated by eigen vectors of the unphosphorylated protein (left) for mode 8 and phosphorylated protein (right) for mode 10. **(E) (i)** Overlap of the first ten non-zero normal modes of phosphorylated and unphosphorylated forms of Lactoperoxidase. **(ii)** Backbone fluctuations indicated by eigen vectors of the unphosphorylated protein (left) and phosphorylated protein (right) for mode 4. * denotes the investigated mode, and thick arrows indicate major collective motions.

Low-frequency global modes derived from Normal Mode Analysis (NMA) have emerged as crucial indicators of biologically relevant dynamics in proteins. They provide insightful information on how these biomolecules function. Previous studies have underscored the significance of these modes in governing enzymatic activity, substrate binding, and conformational changes. For instance, Marcos et al. (2011) demonstrated that enzymes within the amino acid kinase family acquire new modes of motion upon oligomerization [17]. This structural rearrangement plays a pivotal role in regulating substrate binding, highlighting the significance of low-frequency modes in enzymatic function. In a different study, Oliwa and Shen (2015) attempted to model conformational changes that occur upon binding by reevaluating and reordering normal modes [18]. This approach aimed to capture the dynamic rearrangements associated with ligand recognition and binding. Furthermore, investigations have revealed that the low-frequency global modes of unbound proteins are perturbed upon interaction with their binding partners [9]. This phenomenon sheds light on the influence of binding events on the conformational dynamics of proteins.

Building upon these findings, it becomes intriguing to explore the impact of post-translational modifications, such as phosphorylation, on low-frequency global modes. Deciphering whether phosphorylation alters the low-frequency modes of proteins and how these modifications relate to functional dynamics will be crucial to understanding the complex mechanisms that underlie cellular functions. To address this question, we conducted a comprehensive analysis to determine whether phosphorylation introduces new low-frequency motions or preserves the inherent dynamics of proteins. Our approach involved analysing the similarities and differences between the modes of motion accessible to a protein in its phosphorylated and unphosphorylated forms. This was done by quantifying the overlap between the top ten low-frequency global motions obtained from NMA. The overlap value serves as a robust indicator of the similarity between modes in terms of their frequency, shape, and size. A smaller overlap value signifies a larger difference in the two modes of motion. The low-frequency global modes showed little overlap in about 71% of the cases, in our study, suggesting a substantial shift in mode preference and order upon phosphorylation. A shift in mode preference implies that a mode ‘m,’ present as a low-frequency mode in one form, manifests as the same or a similar mode (defined by a high overlap value) in the other form but with a modified frequency. This finding suggests that while some modes of motion are preserved between the phosphorylated and unphosphorylated forms, their frequency, size, and shape undergo substantial changes, as evidenced by the reordering of normal modes. The introduction of a phosphate moiety not only alters the low-frequency modes of proteins but also imparts a distinct character to their functional dynamics. We examined several case examples to provide concrete evidence of the impact of phosphorylation on low-frequency global modes of proteins. One such case involved Protein-arginine-phosphatase (PAP), where we observed that the mode order was retained for the first low-frequency mode, as indicated by high overlap value (**Figure 4C, i**). We observed that the major collective motions of the phosphorylated and unphosphorylated forms were alike, based on the eigen vectors representing the corresponding backbone fluctuation for mode 1 (**Figure 4C, ii**). Another case study focused on Cutinase 1, where we observed a clear instance of mode reordering. Specifically, the eighth mode of the unphosphorylated form manifested as the tenth mode of the phosphorylated protein, as indicated by high overlap values (**Figure 4D, i**). To provide further insight, we examined the corresponding eigen vectors for backbone fluctuation in mode 8 of the unphosphorylated protein and mode 10 of the phosphorylated protein, showcasing a notable degree of comparability (**Figure 4D, ii**). Additionally, we investigated the case of Lactoperoxidase, where we found that the global mode 4 of the unphosphorylated form was not preserved in the phosphorylated form (**Figure 4E, i**). This evidence was supported by the eigen vectors for backbone fluctuation in mode 4 of the unphosphorylated and phosphorylated forms. Notably, the low-frequency motions observed in the unphosphorylated form are lost, and new collective motions are introduced post-phosphorylation, as evident in the phosphorylated protein (**Figure 4E, ii**). The aforementioned case studies collectively provide insight on the diverse effects of phosphorylation on low-frequency global modes-ranging from preserved mode characteristics in PAP to mode reordering in Cutinase 1 and the introduction of new collective motions in Lactoperoxidase.

These observations unveil a fascinating duality in the impact of phosphorylation on protein dynamics. Firstly, the stability in residue motion correlation despite phosphorylation implies that the local interactions and coordinated movements between residues remain relatively unchanged. This suggests a certain robustness in the residue-level dynamics, indicating that phosphorylation may not disrupt the inherent correlations between residues. On the other hand, the perturbation of low-frequency global modes in the unphosphorylated form following phosphorylation reveals a nuanced and intricate alteration in the collective motions of the protein. While the residue-level interactions may remain stable, the larger-scale, global modes of motion are distinctly influenced by phosphorylation. This implies that the post-translational modification introduces changes in the collective, low-frequency dynamics of the protein, potentially impacting its overall structural and functional behaviour. In essence, this duality in the observed effects of phosphorylation highlights the complexity of its influence on protein dynamics.

### Observable differences in the total accessible surface post-phosphorylation are particularly accentuated in the vicinity of the phosphorylation site

The total accessible surface area (ASA) of a protein refers to the combined surface area of all the atoms in the protein that are accessible to the surrounding solvent molecules. It is a measure of the protein’s exposed surface area and can help in understanding the protein’s interactions with other molecules, such as ligands, substrates, or other proteins. Changes in ASA can have significant implications for the protein’s biological activity and its interactions within cellular environments. We sought to better understand the impact of phosphorylation on the total accessible surface area (ASA) of proteins, focusing on identifying specific regions that undergo the most significant changes. We used NetPhos 3.1, a machine learning model trained on a diverse set of experimentally verified phosphorylation events to calculate the ASA values for both phosphorylated and unphosphorylated protein states within the SSP dataset (refer Methods). The exposed and buried residues were distinguished by applying a threshold of 25% relative accessible surface area (RASA). We separately computed the total ASA across distinct residue categories for all protein pairs in the SSP dataset - *Phosphosite (Seq), Non-Phosphosite (Seq), Phosphosite (Str), Near-Phosphosite (Str),* and *Away from Phosphosite (Str).* By comparing the ASA values before and after phosphorylation, we were able to determine the percentage change in the total ASA for the entire protein, as well as for specific regions of interest.

Our findings revealed that 54% of the proteins exhibited a change in the total accessible surface area following phosphorylation. Among these proteins, 25% displayed an increase in the total ASA, while 29% showed a reduction (**Figure 5A**). Notably, 21% of proteins exhibited an augmented number of buried residues coupled with a decrease in exposed residues, whereas 25% displayed an increase in exposed residues and a corresponding decrease in buried residues (**Figure 5B**). Zooming in on specific regions, we observed alterations at the *Phosphosite (Str)* region, where approximately 46% of proteins demonstrated a change in total ASA. Within this category, 17% experienced an increase, while 29% exhibited a decrease in total ASA post-phosphorylation (**Figure 5C, i**). In the *Near-Phosphosite (Str)* region, we noted a substantial 54% showing a percentage difference in total ASA, with a noteworthy 42% of proteins displaying an increase and only 12% showing a reduction (**Figure 5C, ii**). This region exhibited a pronounced skew, signifying a substantial disparity between phosphorylated and unphosphorylated states. In the *Away from Phosphosite (Str)* region, we observed that 54% of the proteins exhibited a percentage difference in total ASA, with 21% showing an increase and 33% showing a decrease (**Figure 5C, iii**). In the *Phosphosite (Seq)* region, approximately 46% of proteins displayed a change, with 21% experiencing an increase and 25% demonstrating a decrease (**Figure 5D, i**). Finally, in the *Non-Phosphosite (Seq)* region, 50% of proteins showcased a difference, with 21% indicating an increase and 29% showing a decrease (**Figure 5D, ii**).

**Figure 5:**
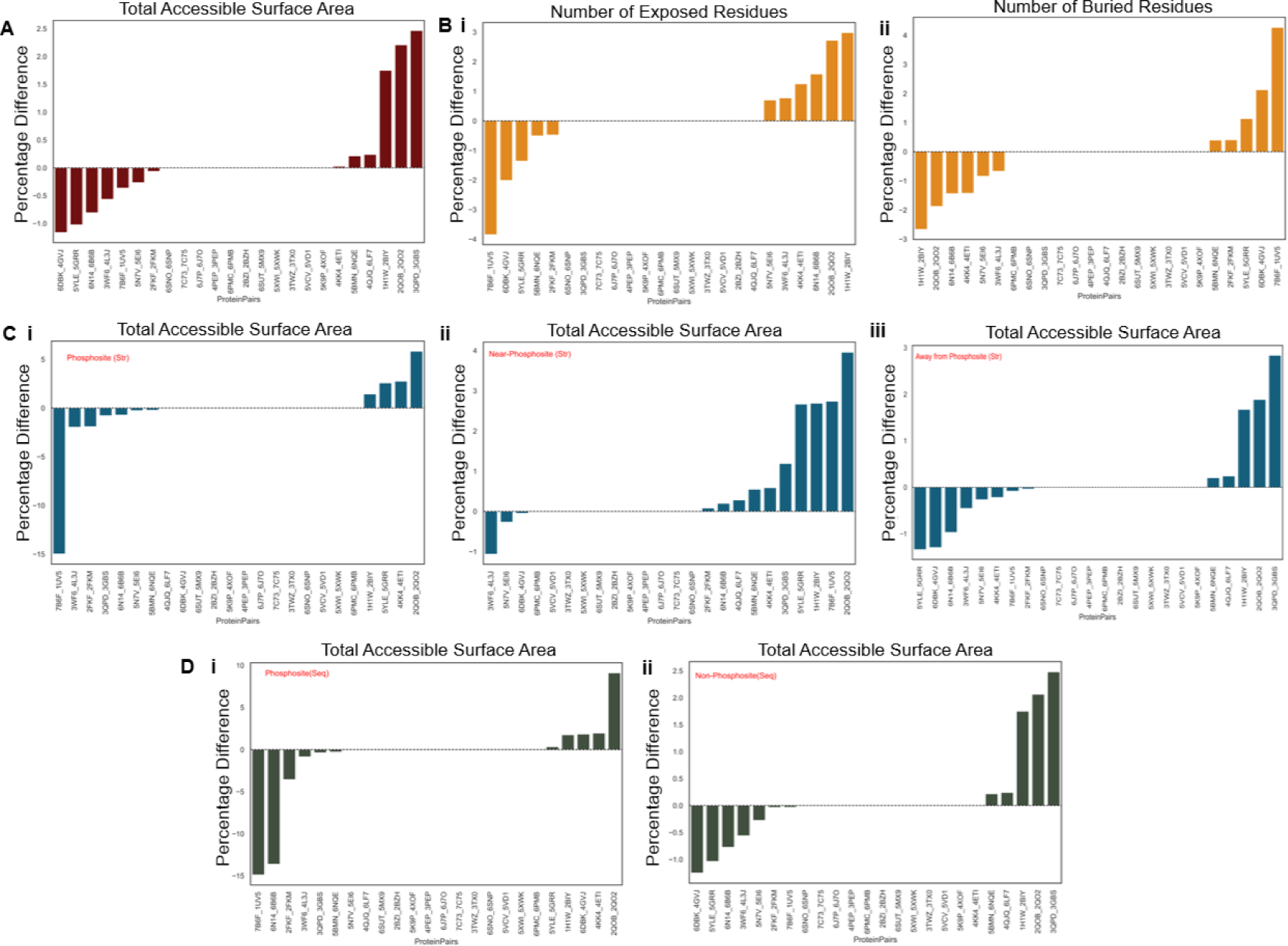
ASA Analyses. **(A)** Percentage change in the total accessible surface area for all proteins. **(B)** Percentage difference (i) in the number of exposed residues post-phosphorylation (ii) in the number of buried residues post-phosphorylation. **(C)** Percentage change in the total accessible surface area considering all (i) Phosphosite (Str) residues (ii) Near-Phosphosite (Str) residues (iii) Away from Phosphosite (Str) residues. **(D)** Percentage change in the total accessible surface area: (i) Phosphosite (Seq) region (ii) Non-Phosphosite (Seq) region. Protein pairs above the black line indicate an increase in surface area post phosphorylation, while protein pairs below the line show a reduction.

By altering the surface accessibility of residues, phosphorylation can regulate protein-protein interactions, signaling pathways, and cellular processes. Based on our analysis, phosphorylation has a notable impact on the accessible surface area (ASA) of proteins. The observed variations in accessible surface area (ASA) post-phosphorylation provide valuable insights into the dynamic nature of protein modifications and their impact on protein structure and function. The increase in total ASA for a subset of proteins suggests that phosphorylation can lead to a more exposed and accessible surface. This phenomenon may be attributed to conformational changes triggered by phosphorylation, exposing previously buried residues to the solvent, potentially facilitating interactions with other molecules or proteins. Conversely, proteins that demonstrated a reduction in total ASA post-phosphorylation imply a more compact or shielded structure. This could be indicative of phosphorylation-induced structural rearrangements that result in a tighter packing of residues or a shielding effect on certain regions of the protein, potentially protecting them from further modifications or interactions. The region-specific differences, particularly in the Near-Phosphosite (Str) category, highlight the intricate and context-dependent nature of phosphorylation effects. The substantial skew in this region, with a majority of proteins showing an increase in total ASA, implies a pronounced impact of phosphorylation in creating more exposed surfaces in proximity to phosphorylation sites, indicating its crucial role in the structural and functional consequences of phosphorylation.

### Phosphorylation at different protein sites induces variable structural and dynamics changes, potentially impacting function: A case study on Polyubiquitin-B

Polyubiquitin B is a protein that plays a crucial role in protein degradation through the ubiquitin-proteasome system. It can exist in different forms: covalently attached to another protein or free (unanchored). When covalently bound, it can be conjugated to target proteins as a monomer (monoubiquitin) or as a polymer linked via different lysine residues of the ubiquitin (polyubiquitin chains). The functional diversity of polyubiquitin chains is contingent upon the specific Lys residue to which ubiquitin is linked [19]. For example, Lys-6-linked chains may be involved in DNA repair, Lys-11-linked chains are associated with endoplasmic reticulum-associated degradation (ERAD) and cell-cycle regulation, Lys-29-linked chains play a role in proteotoxic stress response and cell cycle [20], Lys-33-linked chains are involved in kinase modification, Lys-48-linked chains are crucial for protein degradation via the proteasome, and Lys-63-linked chains participate in endocytosis, DNA-damage responses, and activation of the transcription factor NF-kappa-B. When existing as free entities, unanchored polyubiquitin molecules serve distinct roles, such as activating protein kinases and participating in signaling pathways. Phosphorylation of polyubiquitin B can occur on different residues, leading to diverse functional outcomes. The phosphorylation of specific residues influences the protein’s activity and its interactions with other molecules. For instance, phosphorylation on certain residues may modulate polyubiquitin B’s ability to bind to RNA molecules. Disruption of RNA binding can significantly impact gene expression regulation and other cellular processes. Additionally, phosphorylation at Ser-65 has been shown to activate parkin (PRKN), triggering mitophagy [21]. It has also been observed that Ser-65 phosphorylation affects the discharging of E2 enzymes during the formation of polyubiquitin chains and can impact deubiquitination by enzymes like USP30 [22]. However, it is important to note that the current literature limits comprehensive insights into the distinct functional implications of phosphorylation on different residues, emphasizing the need for further research to elucidate the nuanced effects of residue-specific phosphorylation on polyubiquitin B.

In this study, we analysed two phosphorylated forms of polyubiquitin B: one phosphorylated at serine residue at position 20 (PDB ID: 5K9P) (**Figure 6A, left**) and the other phosphorylated at threonine residue at position 12 (PDB ID: 5NVG) (**Figure 6B, left**), comparing them with the unphosphorylated polyubiquitin B structure (PDB ID: 4XOF). To assess the structure-dynamics changes, we performed structural alignments of the phosphorylated forms with the unphosphorylated form and computed the Calpha deviation and absolute difference of normalized square fluctuations for the aligned residues. We observed that phosphorylation perturbed the low-frequency global modes of the protein, evident from the low overlap values and caused reordering of some modes. Interestingly, the perturbations observed in the Ser20 phosphorylated protein (**Figure 6A, right**) were not necessarily mirrored in the Thr12 phosphorylated protein, underscoring phosphorylated-residue-specific effects (**Figure 6B, right**). For instance, the low-frequency global motions associated with mode 10 of the unphosphorylated form were manifested in mode 7 for Ser20 phosphorylated polyubiquitin but were largely retained in the same mode for Thr12 phosphorylated protein. Similarly, mode 1 motions of the unphosphorylated protein were reordered to mode 2 in Thr12 phosphorylated state but were lost in Ser20 phosphorylated form. The evaluation of Calpha deviation patterns in both Ser20 (**Figure 6C, i**) and Thr12 (**Figure 6C, ii**) phosphorylated proteins compared to the unphosphorylated form unveiled distinct behaviours. While both phosphorylated forms exhibited significant deviation in the β1-β2 loop, the Thr12 phosphorylated form displayed additional significant deviations in regions such as C-α1, α2, β3, β3-β4 loop, and β4. Examining the lysine residues crucial for forming iso-peptide bonds revealed differing degrees of deviations in both Ser20 and Thr12 phosphorylated forms, emphasizing phosphorylated-residue-specific impacts on polyubiquitin chain formation and protein functionality. The patterns of fluctuation difference for Ser20 (**Figure 6D, i**) and Thr12 (**Figure 6D, ii**) phosphorylation proteins compared to the unphosphorylated form were found to be different for each phosphorylated protein. Once again, the β1-β2 region showed significant differences in fluctuation in both cases. In the Ser20 phosphorylated case, the β3-β4 loop also showed significant fluctuation difference, but this was not observed in the Thr12 phosphorylated protein. Significance was attributed only if the difference exceeded the standard deviation from the mean of the fluctuation difference in the control dataset. Furthermore, investigating the global motions for mode 9 of both phosphorylated forms, and the unphosphorylated form revealed nuanced differences (**Figure 6E and Supplementary Movie 1**). Mode 9 was chosen for analysis due to its moderate overlap with the unphosphorylated form in Thr12 phosphorylation and low overlap in Ser20 phosphorylation. This allowed us to compare the low-frequency global motion seen in the unphosphorylated form, which was not retained in the Ser20 phosphorylated form but was marginally preserved in the Thr12 phosphorylated form. Notably, the β1-β2 loop consistently exhibited significant fluctuations in both phosphorylated forms compared to the unphosphorylated form, with greater magnitudes in the phosphorylated states. In the Ser20 phosphorylated case, significant differences in the fluctuations of the β3-β4 loop were observed compared to the unphosphorylated protein, a feature less pronounced in the Thr12 phosphorylated case (as visualized by the length of eigen vectors).

**Figure 6:**
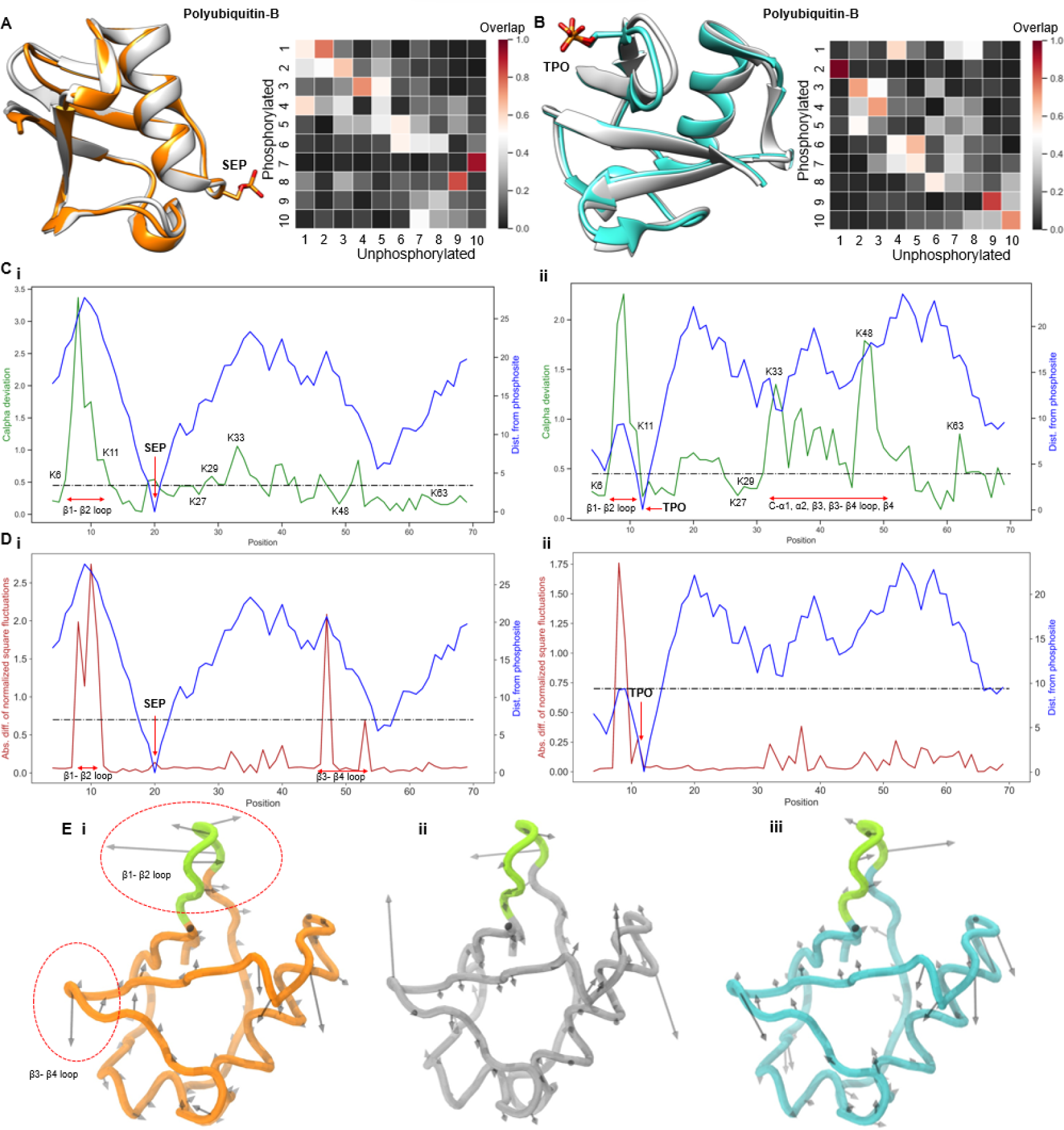
Understanding Structure-Dynamic Alterations in Polyubiquitin-B. **(A)** Left: Superimposed and aligned structures of Ser 20 phosphorylated (PDB ID: 5K9P, coloured orange) and unphosphorylated (PDB ID: 4XOF, coloured silver) forms. Right: Overlap of the first ten non-zero normal modes of both forms. **(B)** Left: Superimposed and aligned structures of Thr 12 phosphorylated (PDB ID: 5NVG, coloured cyan) and unphosphorylated (PDB ID: 4XOF, coloured silver) forms. Right: Overlap of the first ten non-zero normal modes of both forms. **(C)** Cα-deviations between (i) Ser 20 phosphorylated and unphosphorylated forms (ii) between Thr 12 phosphorylated and unphosphorylated forms for all aligned residues, denoted by a green line. **(D)** (i) Absolute difference between normalized square fluctuations of (i) Ser 20 phosphorylated and unphosphorylated forms (ii) Thr 12 phosphorylated and unphosphorylated forms, for aligned residues, denoted by a maroon line. The blue line represents the distance of each residue from the phosphorylation site. The black line indicates the significance cutoff. Regions/residues of interest are marked in the plots. **(E)** Backbone fluctuations indicated by eigenvectors of mode 9 for Ser 20 phosphorylated form (left), unphosphorylated form (middle), and Thr 12 phosphorylated form (right).

Overall, this case study highlights that the type and position of the phosphorylated residue intricately influence the structural dynamics of polyubiquitin B, potentially leading to divergent functional outcomes. The observed variations in global modes, local structural deviations, and fluctuations provide valuable insights into the residue-specific impact of phosphorylation on the structural and dynamic properties of polyubiquitin B.

### Phosphorylation within a kinase’s activation loop brings about functional changes by influencing its structure and dynamics: A case study on Glycogen Synthase Kinase-3 Beta

Glycogen Synthase Kinase-3 Beta (GSK-3β) is a serine/threonine protein kinase that plays a crucial role in various cellular processes, including glycogen metabolism, cell proliferation, differentiation, and apoptosis. One important aspect of GSK-3β regulation is its phosphorylation at Tyr216, which activates the kinase. This phosphorylation event is often associated with increased GSK-3β activity. Activated GSK-3β, in turn, phosphorylates a range of substrate proteins, exerting regulatory control over diverse cellular functions[23, 24]. The phosphorylation of GSK-3β at Tyrosine 216 has been implicated in several biological processes. For instance, it has been shown to be involved in mechanisms of neuronal survival [25]. Additionally, the phosphorylation of GSK-3β at Tyrosine 216 has been linked to the regulation of tau protein, which is associated with neurodegenerative diseases like Alzheimer’s disease [26, 27]. Abnormal GSK-3β activity can contribute to pathological processes associated with these diseases. A comprehensive understanding of GSK-3β’s role in cellular signaling necessitates a detailed exploration of its phosphorylation patterns.

We conducted a comparative conformational analysis of GSK-3β in its Tyr216 phosphorylated and unphosphorylated forms. The two forms exhibited a Cα-RMSD of 1.62 Å, highlighting substantial structural disparities (**Figure 7A**). A detailed examination of the aligned residues revealed significant backbone deviations in the N-terminal lobe, particularly the β3-αC loop and specific regions of the C-terminal lobe (**Figure 7B**). Furthermore, the N-terminal lobe displayed a higher difference in flexibility, particularly in the Glycine-rich loop compared to the C-terminal lobe. This observation aligns with the characteristic motions observed in active kinases (**Figure 7C**). Importantly, the regions exhibiting significant flexibility were found to be distant from the phosphorylation site, suggesting long-range dynamic changes resulting from phosphorylation. Additionally, we investigated the preservation of low-frequency global modes and found that only a limited subset of modes was preserved post-phosphorylation, with a majority being lost and a few modes undergoing re-ordering (**Figure 7D**). A focused investigation into mode 5 motions demonstrated distinct differences between the two forms. The phosphorylated kinase exhibited characteristic lobe-opening and closing motions, reminiscent of active kinases, a motion absent in the unphosphorylated form. Moreover, the β3-αC loop in the unphosphorylated form displayed substantial fluctuations, contributing to significant differences in fluctuation dynamics in this region (**Figure 7E and Supplementary Movie 2**). Our findings highlight substantial conformational disparities between the phosphorylated and unphosphorylated forms of GSK-3β, revealing specific structural and dynamic alterations induced by phosphorylation, including long-range effects and alterations in low-frequency global modes.

**Figure 7:**
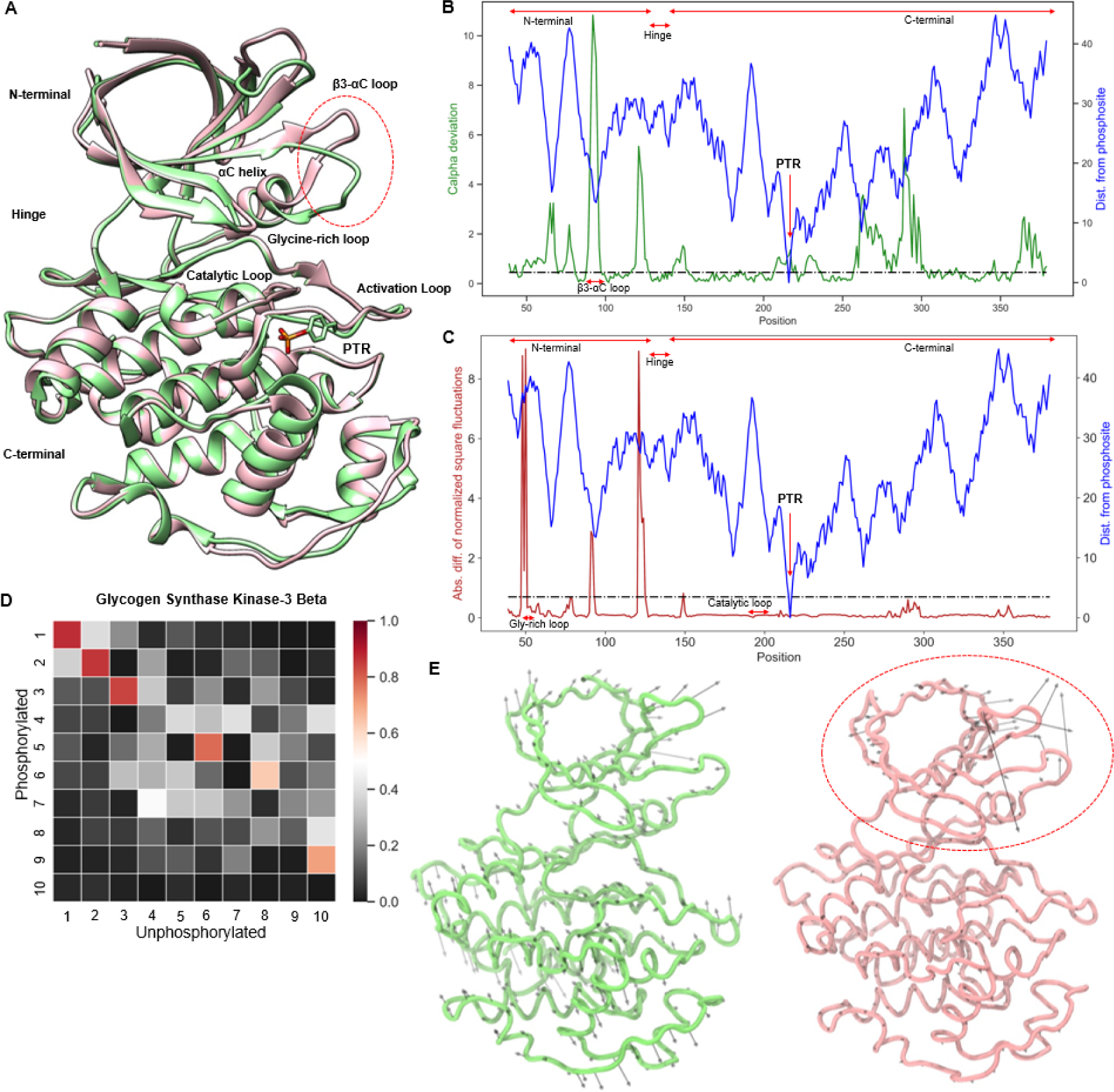
Phosphorylation Induces Changes in Glycogen Synthase Kinase-3 Beta. **(A)** Superimposed and aligned structures of Tyr 216 phosphorylated (PDB ID: 7B6F, coloured green) and unphosphorylated (PDB ID: 1UV5, coloured pink) forms of GSK -3β. Important regions of the kinase are marked, and the red circle indicates the region with significant structural disparities. **(B)** C-alpha deviation between phosphorylated and unphosphorylated forms for all aligned residues, denoted by a green line. **(C)** Absolute difference between normalized square fluctuations of phosphorylated and unphosphorylated forms for aligned residues, denoted by a maroon line. The blue line represents the distance of each residue from the phosphorylation site. The black line indicates the significance cutoff. Regions/residues of interest are marked in the plots. **(D)** Overlap of the first ten non-zero normal modes of both phosphorylated and unphosphorylated forms. **(E)** Backbone fluctuations indicated by eigenvectors of mode 5 for the phosphorylated form (left) and unphosphorylated form (right).

## DISCUSSION

Our study delves into the intricate landscape of phosphorylation-induced conformational dynamics, providing a detailed investigation of the alterations in protein structure, dynamics, and surface accessibility. The nuanced findings shed light on the multifaceted impact of single-site phosphorylation, uncovering both local and global effects that contribute to the regulatory tapestry of cellular processes. One of the key observations from our study is the distinctive conformational changes that proteins undergo upon phosphorylation. The structural alterations extend far beyond the phosphorylation site, with more pronounced changes occurring distally. This emphasizes the ripple effect of phosphorylation on protein structure, indicating a sophisticated network of conformational adjustments that intricately shape the functional properties of proteins. Our exploration of phosphorylation-induced dynamic shifts in protein residues through Normal Mode Analysis (NMA) provided nuanced insights into alterations in vibrational modes and flexibility. The observed significant differences in normalized square fluctuations between phosphorylated and unphosphorylated forms indicate a post-phosphorylation augmentation in the dynamic motion of residues. The robustness of these findings was confirmed through consistent observations across different distance cutoffs, reinforcing the reliability of our results. Contrary to the dynamic shifts observed, residue motion correlation remained largely stable post-phosphorylation, indicating robustness in the local interactions and coordinated movements between residues. However, an intricate duality emerged when exploring low-frequency global modes. The low overlap observed in a majority of cases suggests a substantial shift in mode preference and order upon phosphorylation. This emphasizes that while local interactions may persist, the larger-scale, global modes of motion are distinctly influenced by phosphorylation, showcasing a nuanced alteration in the collective dynamics of the protein. The investigation into the total accessible surface area (ASA) post-phosphorylation revealed diverse effects on protein exposure and interaction capabilities. A substantial portion of proteins exhibited changes in total ASA, with some showcasing an increase and others a reduction. The region-specific differences, especially in the Near-Phosphosite (Str) category, underscore the localized impact of phosphorylation in creating more exposed surfaces in proximity to phosphorylation sites. The case study on Polyubiquitin B offers a gripping insight into the residue-specific phosphorylation effects on the structural dynamics, providing a nuanced understanding of the molecular mechanisms underlying its functionality. This case underscores the need for future research to unravel the distinct functional implications of phosphorylation on different residues and at different locations, contributing to a more comprehensive understanding of the residue-position-specificity. The case study on Glycogen Synthase Kinase-3 Beta (GSK-3β) provides a detailed exploration of how phosphorylation within the activation loop at Tyr216 influences the structure and dynamics of the kinase, thereby impacting its functionality in various cellular processes. The long-range effects, preservation of specific global modes, and residue-specific fluctuation dynamics underscore the intricate relationship between phosphorylation and kinase activity. Future research endeavours can build upon these findings, further elucidating the molecular mechanisms that govern GSK-3β regulation and its implications for cellular processes and disease pathology. A primary limitation of our research is the availability of suitable crystal structures in both phosphorylated and unphosphorylated forms in the Protein Data Bank (PDB). The stringent filtering criteria applied, necessitated by the nature of our study, further reduced the number of eligible protein pairs. This dual constraint, stemming first from the limited availability of relevant structures and then from our rigorous filtration process, restricted the size and diversity of our dataset. Consequently, the outcomes and generalizability of our findings may be influenced by this inherent limitation in the selection of proteins for analysis. Despite these constraints, our study provides valuable insights into the structural and dynamic effects of protein phosphorylation, paving the way for future investigations with an expanded and more diverse dataset.

In conclusion, our study provides valuable insights into the conformational, dynamic, and surface accessibility alterations induced by phosphorylation. By elucidating the intricacies of phosphorylation-induced structure-dynamics-accessibility changes, we contribute to a more comprehensive understanding of the regulatory mechanisms underlying cellular processes. This knowledge can pave the way for the development of targeted therapeutic strategies aimed at modulating phosphorylation events and their associated cellular signaling networks.

## MATERIALS AND METHODS

### Dataset Preparation

#### SSP dataset for structural comparison

A dataset comprising 24 non-redundant protein pairs, constituting a total of 48 (24*2) protein structures, was systematically curated from the RCSB Protein Data Bank (**Table S1**). Each pair consisted of both phosphorylated (in Serine/Threonine or Tyrosine, denoted as SEP/TPO/PTR) and unphosphorylated forms. The data curation process involved applying stringent filtering conditions:

1. All protein pairs must have 3-D crystal structures available in both phosphorylated and unphosphorylated states, with no missing residues at or near the phosphorylated residue.
2. The resolution of the structures should be better than 3 Å.
3. Both phosphorylated and unphosphorylated forms must share the same UniProt identifier.
4. The oligomeric state of both the forms should be identical (here, AU=1, BA=1), as determined by examining PDB biological unit information.
5. Both forms should either have similar ligands or no ligands bound. This condition was imposed to minimize and mitigate biases arising due to the presence of ligand(s).
6. Exclusion of proteins with disordered regions.
7. Both phosphorylated and unphosphorylated forms should either have no mutation or identical mutations.
8. The phosphate moiety should be present as a modified residue, distinct from the classification as a ligand.

These rigorous criteria were implemented to ensure that any observed differences between the phosphorylated and unphosphorylated forms are solely attributed to the addition of a phosphate moiety.

#### SSP dataset for dynamics analysis

To understand the nuanced dynamics and flexibility changes within protein structures upon phosphorylation, the initial dataset underwent additional refinement. Instances with missing residues in either the phosphorylated or unphosphorylated state were excluded. Enforcement of this filtering condition aimed to preclude potential bias stemming from the necessity to model missing regions in the protein structure. Consequently, 17 protein pairs, totalling 34 distinct protein structures, satisfying the refined criteria were identified and selected for subsequent analyses.

#### Control dataset

The control group consisted of phosphorylated protein structures from representative protein pairs in the SSP dataset, resolved under varied crystallographic conditions. These structures were further filtered to ensure the presence of single chain in both the asymmetric unit (AU) and biological assembly (BA), to maintain congruence with the oligomeric state of protein pairs in the SSP dataset. Care was taken to ascertain the absence of additional biological entities, such as peptides, RNA, or DNA, in both the AU and BA. Further refinement involved applying a resolution cut-off of 3 Å, and exclusion of structures with missing residues. A refined subset comprising 98 structures was thus identified and selected as the control for subsequent analyses. This carefully curated control dataset serves as a reference to elucidate the impact of crystal packing on protein conformation. Importantly, it serves as a baseline, representing background noise, against which significant differences between phosphorylated and unphosphorylated forms can be discerned and quantified.

### Classification of Localized Regions by Integrating Sequence and Structural Perspectives

In this study, protein residues were classified into distinct categories based on two key criteria: sequence and structural proximity.

#### Sequence-Based Classification

The sequence-centric approach involves categorization of protein residues into two groups: “*Phosphosite*” and “*Non-Phosphosite*”. *Phosphosite (Seq)* is defined as heptapeptide spanning from P-3 to P+3, with the ‘P’ representing the phosphorylated residue, while *Non-Phosphosite (Seq)* includes all residues outside this heptapeptide range. This heptapeptide motif, with the phosphorylated residue at the central position (P) and surrounding residues labelled as P-3 to P+3, is chosen due to its extensive use in the literature for kinase substrate identification [28, 29]. This motif aligns with kinase recognition specificity, making it a valuable starting point in studying phosphorylation events.

#### Structural Proximity Classification

The structural classification zooms in on spatial relationships, further dividing localized regions into “*Phosphosite”*, “*Near-Phosphosite”*, and “*Away from Phosphosite*” based on proximity to the phosphorylated residue, employing specific distance criteria. *Phosphosite (Str)* includes residues within a 4.5 Å proximity, determined by an interatomic distance cut-off. The *Near-Phosphosite (Str)* category encompasses residues situated more than 4.5 Å but less than 10 Å away, while the *Away from Phosphosite (Str)* group includes residues positioned more than 10 Å from the phosphorylated residue.

### Structural Analyses

TM-align method was employed to align the structures of phosphorylated and unphosphorylated forms within each protein pair. TM-align, a widely recognized structural alignment algorithm in bioinformatics, facilitates the comparison and alignment of protein structures by emphasizing both sequential and spatial information [30]. The algorithm proves particularly useful for comparing proteins with similar overall folds but potentially different local conformations, a scenario often encountered in studies involving phosphorylated and unphosphorylated forms of proteins.

Following the structural alignment, the backbone root-mean-square deviation (RMSD) was computed as a metric to quantify the dissimilarity between the aligned structures. RMSD is a measure of the average distance between corresponding atoms in two structures. Mathematically, the RMSD between two sets of coordinates (X1, Y1, Z1) and (X2, Y2, Z2) for a set of atoms is calculated as follows:

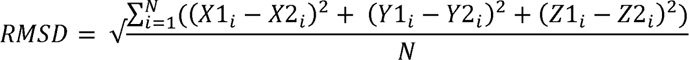

Here, N represents the number of atoms considered in the calculation. A higher RMSD value indicates lower structural similarity, with values exceeding the standard deviation from the mean of RMSDs in the control dataset being considered as statistically significant differences. Additionally, we computed the TM-score, which provides a complementary assessment of structural similarity.

To compute local structural variations, individual alpha carbon (Cα) deviations between equivalent residue positions in both phosphorylated and unphosphorylated forms were calculated. The RMSD between phosphorylated and unphosphorylated proteins was computed for specific regions, including *Phosphosite (Seq), Non-Phosphosite (Seq), Phosphosite (Str), Near-Phosphosite (Str),* and *Away from Phosphosite (Str).* These analyses were conducted for each protein pair in the SSP dataset, providing a comprehensive understanding of localized structural deviations in relation to phosphorylation states.

### Coarse-Grained Dynamics Analysis (Utilizing Normal Mode Analysis with ANM)

Understanding long-timescale protein motions is crucial in deciphering their functional significance, and Normal Mode Analysis (NMA) stands out as a preferred method for such investigations [31–33]. NMA relies on a set of Cartesian coordinates derived from the protein structure and a force field defining inter-atomic interactions. The process involves generating a ‘Hessian’ matrix from the second derivative of potential energy, followed by diagonalization to yield eigenvalues and eigenvectors.

Previous studies have demonstrated that global protein motions, characterized by low-frequency collective movements, play a pivotal role in signifying biologically relevant functions [32, 34]. The computational challenge posed by all-atom NMA, given its extensive calculations, led us to adopt a coarse-grained approach using the Cα-level Anisotropic Network Model (ANM)-based NMA for this study [35]. The choice of Cα-level NMA was grounded in its ability to successfully corroborate with both experimental and molecular dynamics data, providing insights into dynamics over extended timescales [36, 37].

The calculations pertaining to normal mode analyses were executed using the ProDy package [38]. Normal modes were computed for 17 protein pairs in both phosphorylated and unphosphorylated forms, resulting in a total of 34 (17*2) calculations, alongside a control dataset.

For the coarse-grained model, Cα atoms, represented as masses, were connected by elastic springs with identical spring constants if their inter-Cα distance was less than 15 Å, with variations tested using 12 Å and 10 Å cutoffs. Exclusion of contributions from three N-terminal and three C-terminal residues was implemented. Mean squared fluctuations were scaled using z-score normalization, and significance in the differences between normalized square fluctuations of phosphorylated and unphosphorylated forms was determined by comparing them to the standard deviation from the mean of differences in the control dataset. This analysis was conducted for the entire protein and specific regions, including *Phosphosite (Seq), Non-Phosphosite (Seq), Phosphosite (Str), Near-Phosphosite (Str),* and *Away from Phosphosite (Str)*. To quantify dynamics, the Root Mean Squared Difference of Fluctuations (RMSDf) was computed as an equivalent of RMSD, involving the difference between normalized fluctuations of phosphorylated and unphosphorylated forms.

Cross-correlation, indicating the correlation between fluctuations, was computed for all structures. The R_v_ coefficient, a multivariate generalization of Pearson’s coefficient, was employed to quantify the similarity between cross-correlation matrices [39]. Furthermore, to gauge the accessibility of intrinsic motions in phosphorylated form to the unphosphorylated form, the Overlap, representing the inner product of eigenvectors from the 10 lowest frequency modes, was calculated using the Prody package [40, 41]. This provided insight into the extent of shared conformational space between the two forms, shedding light on the protein’s dynamical behaviour.

### Solvent Accessibility Analyses

Solvent accessibility refers to the degree to which a residue is exposed or buried within the protein structure. Solvent accessibility analyses play a crucial role in understanding the structural characteristics of proteins [42]. In this study, a specialized tool NetPhos 3.1 was employed to assess the solvent accessibility of proteins in the SSP dataset. NetPhos 3.1 is a machine learning model that has been trained on a diverse dataset of experimentally verified phosphorylation events [43].

To determine the solvent accessibility of specific residues, NetPhos 3.1 utilizes a sequence-based method. It distinguishes between exposed and buried residues by applying a threshold of 25% relative accessible surface area (RASA). NetPhos 3.1 was chosen for computing solvent accessibility in phosphorylated proteins due to its specialized training on experimental phosphorylation cases, enabling accurate prediction of solvent accessibility for phosphorylated residues and offering insights into their functional and structural implications.

To assess the impact of phosphorylation, we calculated the total accessible surface area (ASA) for both phosphorylated and unphosphorylated forms of the proteins. Equivalent residues in both states were taken into consideration during the analysis. Furthermore, we computed the total accessible surface area for specific regions, including the *Phosphosite (Seq), Non-Phosphosite (Seq), Phosphosite (Str), Near-Phosphosite (Str),* and *Away from Phosphosite (Str)* regions, for each protein pair in the SSP dataset.

To quantify the change in solvent accessibility, we calculated the percentage change in the total accessible surface area as follows:

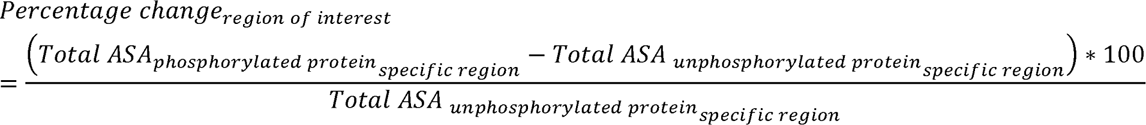

### Statistical Analyses

Statistical significance was assessed using both the 2-sample Kolmogorov-Smirnov (KS) test and the T-Test to scrutinize the distributional disparities between two groups – phosphorylated proteins and unphosphorylated proteins in the SSP dataset. All statistical analyses were carried out in the Python programming environment.

### Visualization

UCSF Chimaera [44] was used for protein structure visualisation, and ".nmd" extension files created using ProDy were used to explore normal modes in Visual Molecular Dynamics (VMD) [45] with the help of the NMWiz (Normal Mode Wizard) plugin. For comparative assessments of normal modes, informative videos were produced using the MovieMaker plugin in VMD, complemented by VideoMach.

## ACKNOWLEDMENTS

Ramanathan Sowdhamini acknowledges funding and support provided by JC Bose Fellowship (JBR/2021/000006) from Science and Engineering Research Board, India and Bioinformatics Centre Grant funded by Department of Biotechnology, India (BT/PR40187/BTIS/137/9/2021). Ramanathan Sowdhamini would also like to thank Institute of Bioinformatics and Applied Biotechnology for the funding through her Mazumdar-Shaw Chair in Computational Biology (IBAB/MSCB/182/2022). Seemadri Subhadarshini is supported by PMRF (Prime Ministers’ Research Fellowship) awarded by Government of India.

## AUTHOR CONTRIBUTIONS

SS conducted research, analysed data, and prepared the initial draft of the manuscript. NS conceptualized the project and provided supervision. HT also provided supervision for the project and reviewed the first draft. RS administered the project, secured research funding, and reviewed the manuscript.

## SUPPLEMENTARY MATERIAL

**Table S1:**
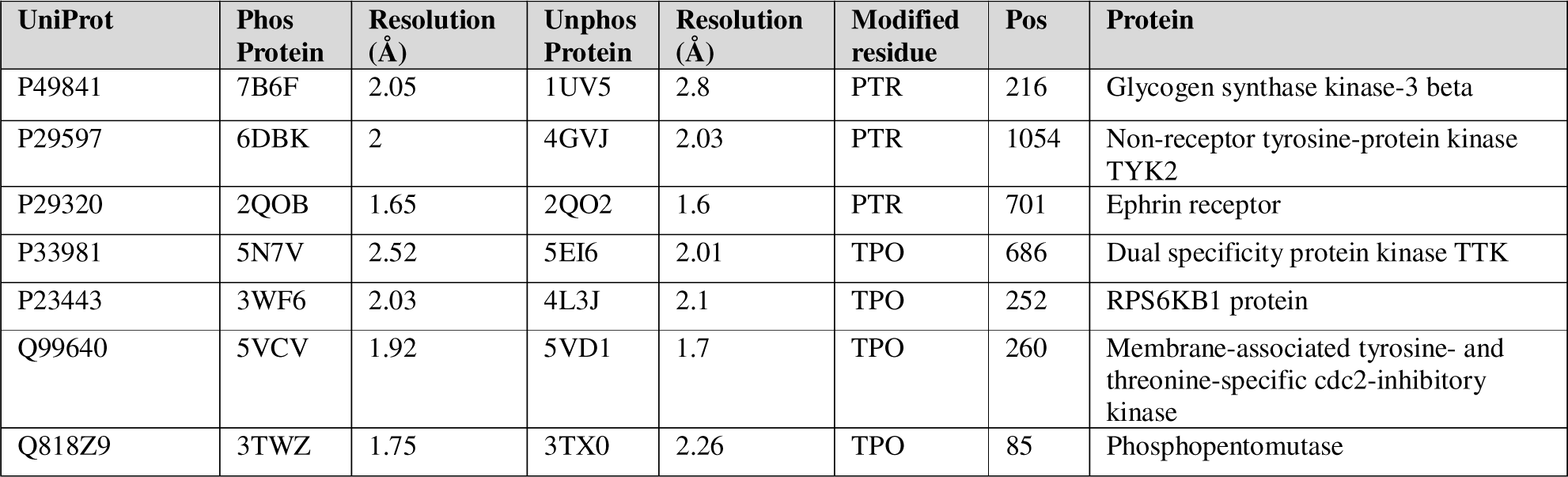
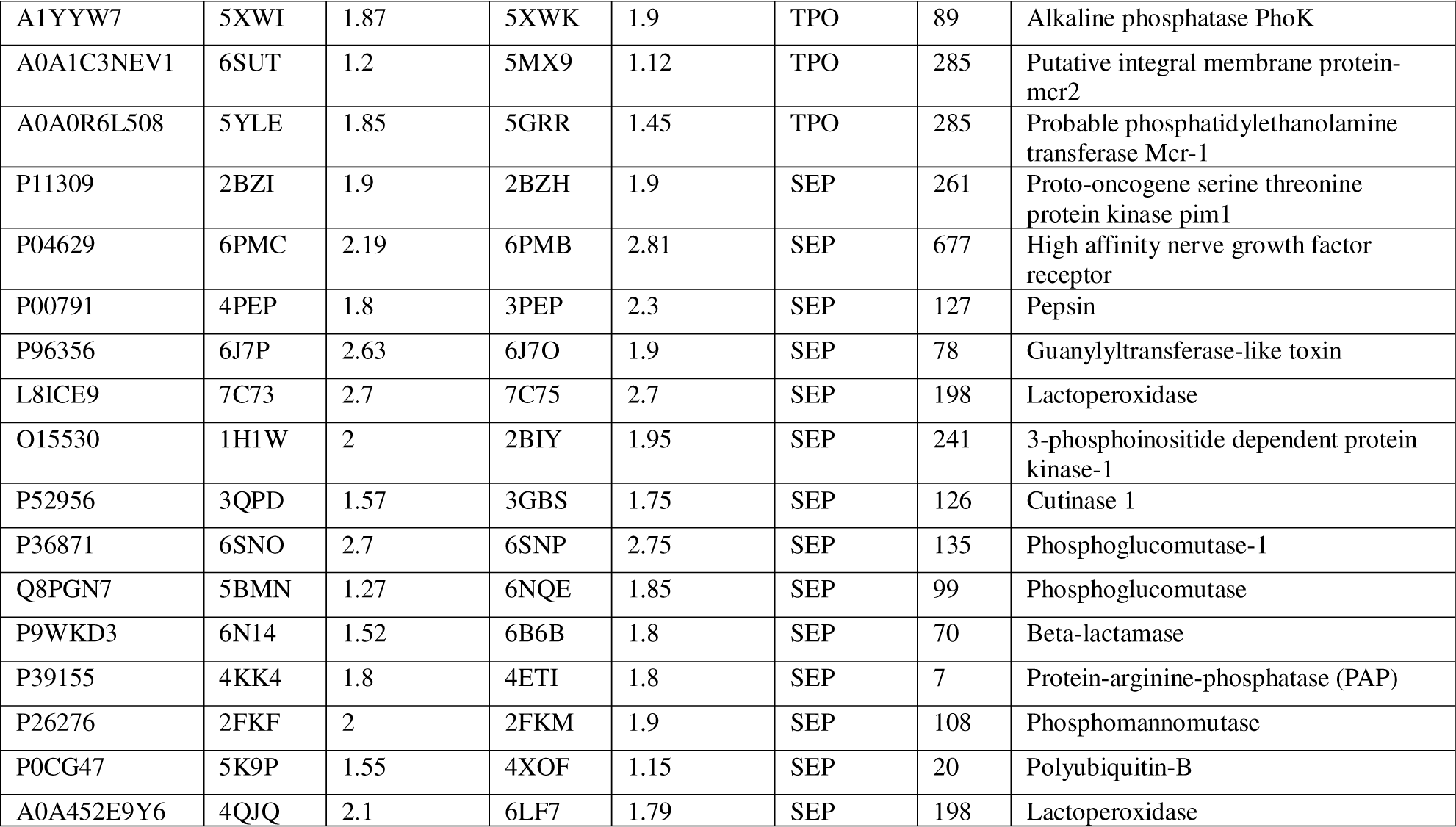
SSP dataset.

**Supplementary Figure 1:**
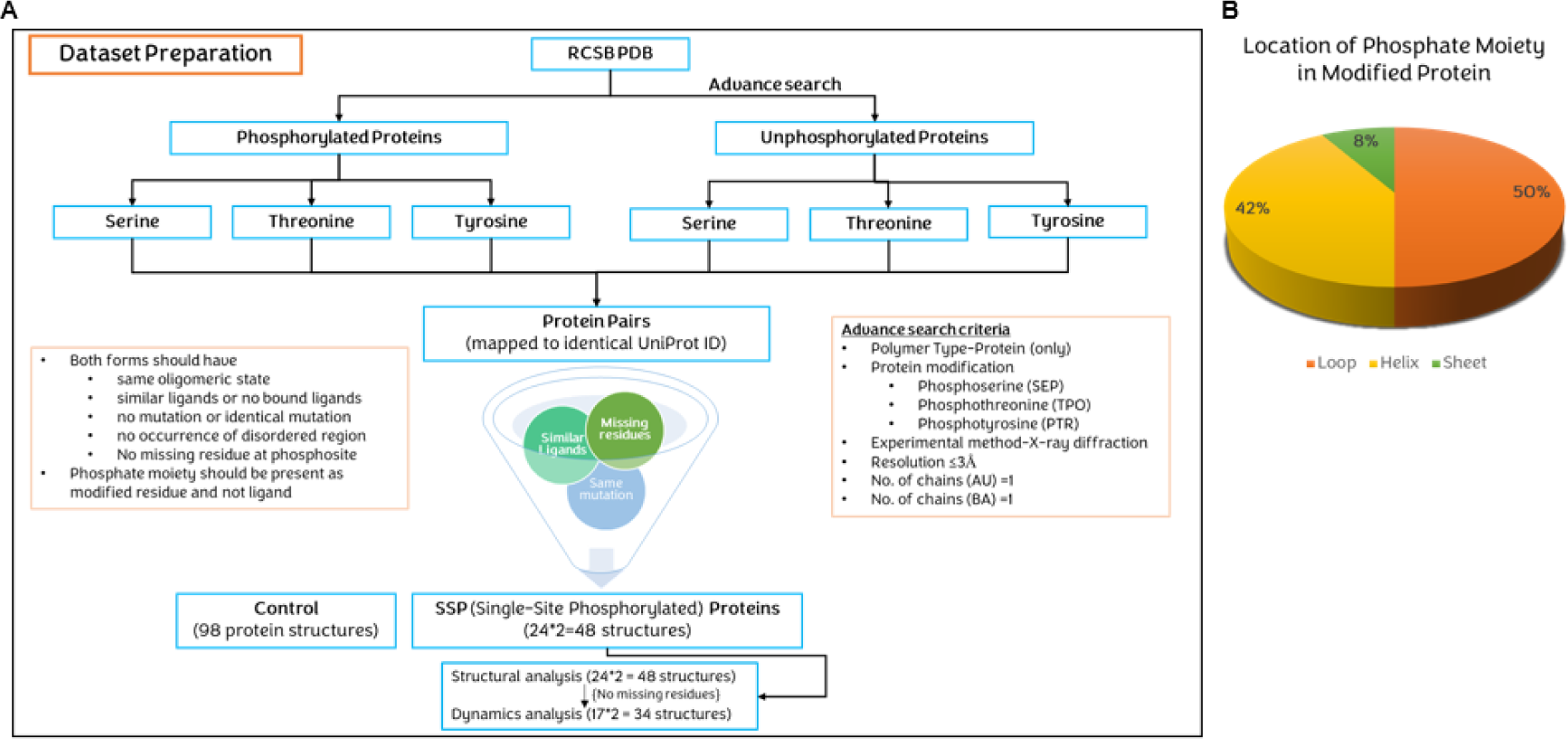
Workflow for SSP Dataset curation and modified residue localization in phosphorylated Proteins. **(A)** Schematic illustration of the dataset preparation workflow for the SSP dataset. (B) Pie chart depicting the specific locations of phosphate moieties in phosphorylated proteins within the SSP dataset.

**Supplementary Figure 2:**
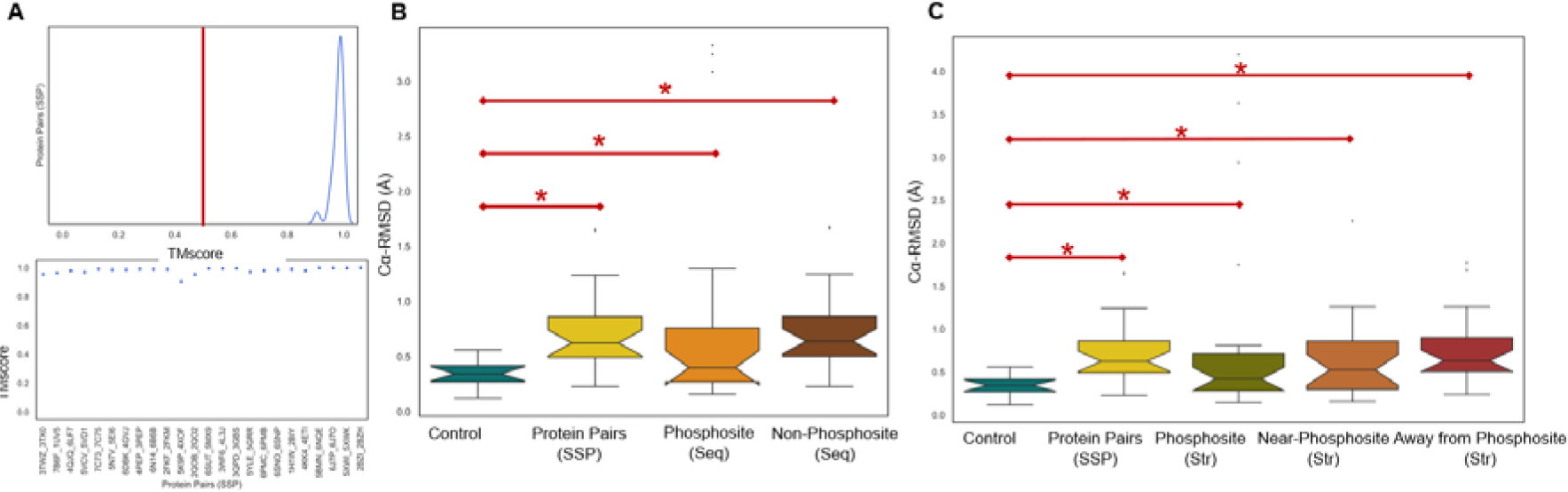
Structural insights into the SSP Dataset (A) Top panel: Kernel Density Estimation (KDE) plot of the distribution of TM scores for all proteins in the SSP dataset. Bottom panel: Scatterplot of TM scores for each protein pair in the SSP dataset. **(B)** Box plots of the distribution of Cα-RMSD (Å) for the Control dataset, Protein Pairs in the SSP dataset, Phosphosite (Seq), and Non-Phosphosite (Seq), with outliers. **(C)** Box plots displaying the distribution of Cα-RMSD (Å) for the Control dataset, Protein Pairs in the SSP dataset, Phosphosite (Str), Near-Phosphosite (Str), and Away from Phosphosite (Str), with outliers. (*) denote significant differences in distributions (evaluated using two-sample KS test and T-test, with a p-value < 0.01).

**Supplementary Figure 3:**
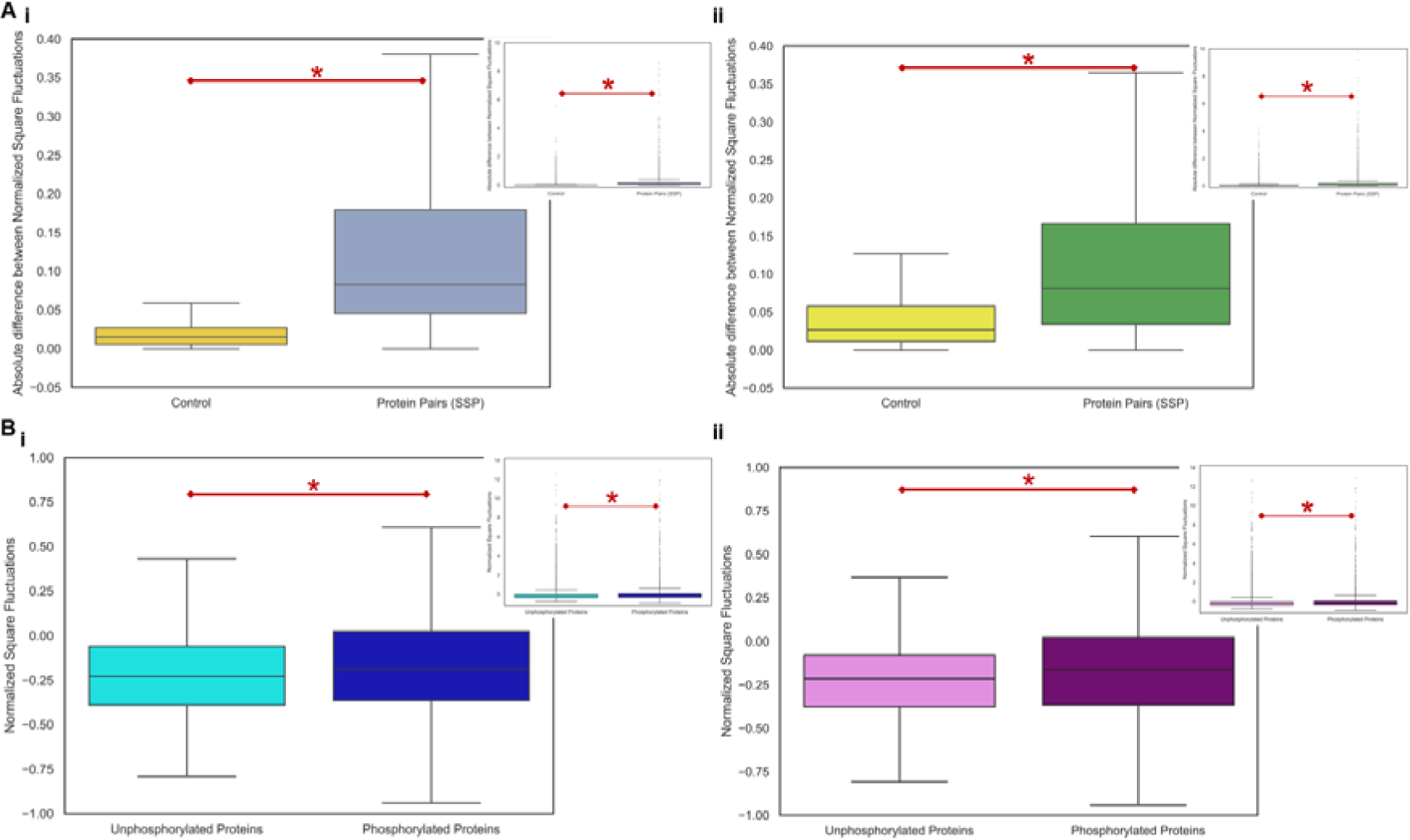
Normal Mode Fluctuations Calculation at 12 Å and 10 Å Cut-off. **(A)** Box plot illustrating the distribution of the absolute difference in normalized square fluctuation of proteins between the control dataset and the phosphorylated/unphosphorylated forms in the SSP dataset at two cut-off values: i) 12 Å cut-off ii) 10 Å cut-off. (B) Box plot displaying the distribution of normalized square fluctuations for all residues in phosphorylated and unphosphorylated proteins at two cut-off values: i) 12 Å cut-off ii) 10 Å cut-off. Inset plot with outliers is included. * indicates significant differences in distributions, determined using a two-sample KS test with a p-value < 0.01.

**Supplementary Figure 4:**
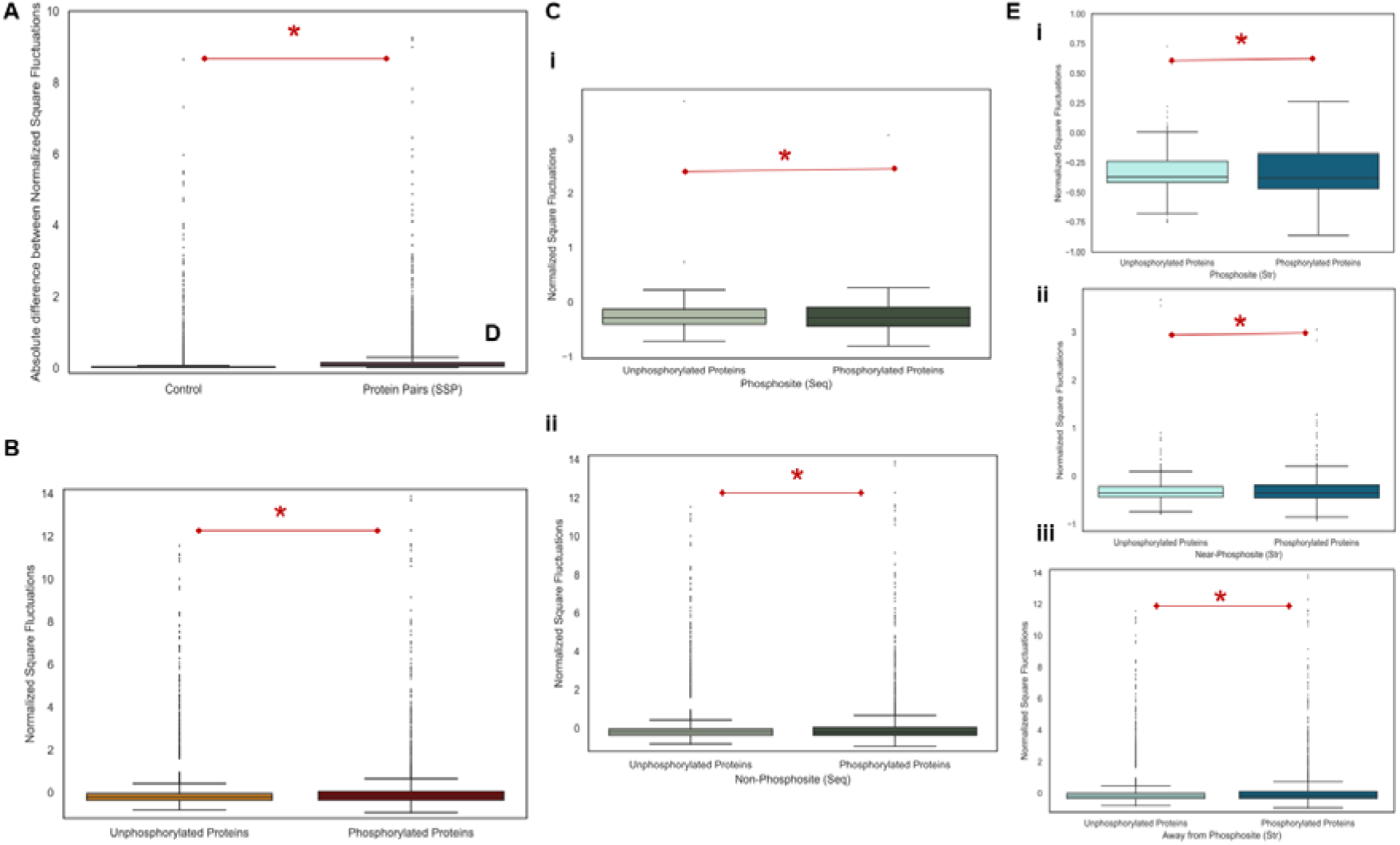
Dynamic/Flexibility Alterations in SSP Dataset. **(A)** Box plots illustrating the distribution of the absolute difference in normalized square fluctuation of proteins between the control dataset and the phosphorylated/unphosphorylated forms in the SSP dataset, including outliers. **(B)** Box plot displaying the distribution of normalized square fluctuations for all residues in phosphorylated and unphosphorylated proteins, including outliers. **(C)** Box plot displaying the distribution of normalized square fluctuations for: i) Phosphosite (Seq) residues ii) Non-Phosphosite (Seq) residues in phosphorylated and unphosphorylated proteins. **(D)** Box plot displaying the distribution of normalized square fluctuations for: **i)** Phosphosite (Str) residues **ii)** Near-Phosphosite (Str) residues **iii)** Away from Phosphosite (Str) residues in phosphorylated and unphosphorylated proteins, including outliers. * indicates significant differences in distributions, determined using a two-sample KS test with a p-value < 0.01.

